# DeepVID v2: Self-Supervised Denoising with Decoupled Spatiotemporal Enhancement for Low-Photon Voltage Imaging

**DOI:** 10.1101/2024.05.16.594448

**Authors:** Chang Liu, Jiayu Lu, Yicun Wu, Xin Ye, Allison M. Ahrens, Jelena Platisa, Vincent A. Pieribone, Jerry L. Chen, Lei Tian

**Affiliations:** Boston University, Department of Biomedical Engineering, Boston, MA 02215, USA; Boston University, Department of Electrical and Computer Engineering, Boston, MA 02215, USA; Boston University, Department of Computer Science, Boston, MA 02215, USA; Neurophotonics Center, Boston University, Boston, MA 02215, USA; Boston University, Department of Biology, Boston, MA 02215, USA; Yale University, Department of Cellular and Molecular Physiology, New Haven, CT 06520, USA; The John B. Pierce Laboratory, New Haven, CT 06519, USA; Yale University, Department of Neuroscience, New Haven, CT 06520, USA

**Keywords:** deep learning, self-supervised denoising, voltage imaging, low photon, microscopy

## Abstract

**Significance:** Voltage imaging is a powerful tool for studying the dynamics of neuronal activities in the brain. However, voltage imaging data are fundamentally corrupted by severe Poisson noise in the low-photon regime, which hinders the accurate extraction of neuronal activities. Self-supervised deep learning denoising methods have shown great potential in addressing the challenges in low-photon voltage imaging without the need for ground truth, but usually suffer from the tradeoff between spatial and temporal performance.

**Aim:** We present DeepVID v2, a novel self-supervised denoising framework with decoupled spatial and temporal enhancement capability to significantly augment low-photon voltage imaging.

**Approach:** DeepVID v2 is built on our original DeepVID framework,^1,2^ which performs frame-based denoising by utilizing a sequence of frames around the central frame targeted for denoising to leverage temporal information and ensure consistency. The network further integrates multiple blind pixels in the central frame to enrich the learning of local spatial information. Additionally, DeepVID v2 introduces a new edge extraction branch to capture fine structural details in order to learn high spatial resolution information.

**Results:** We demonstrate that DeepVID v2 is able to overcome the tradeoff between spatial and temporal performance, and achieve superior denoising capability in resolving both high-resolution spatial structures and rapid temporal neuronal activities. We further show that DeepVID v2 is able to generalize to different imaging conditions, including time-series measurements with various signal-to-noise ratios (SNRs) and in extreme low-photon conditions.

**Conclusions:** Our results underscore DeepVID v2 as a promising tool for enhancing voltage imaging. This framework has the potential to generalize to other low-photon imaging modalities and greatly facilitate the study of neuronal activities in the brain.

## 1 Introduction

Voltage imaging is a powerful tool for studying the dynamics of neuronal activities in the brain. It enables the visualization of the spatiotemporal patterns of membrane potential changes in neurons, which is critical for understanding the underlying mechanisms of brain functions.^3, 4^ Recently, two-photon imaging has also been adapted for voltage imaging, as it provides high spatial resolution and deep tissue penetration.^1, 5, 6^ However, voltage imaging data is often corrupted by strong noise, which hinders the accurate extraction of neuronal activities. The noise in voltage imaging data is mainly attributed to the low photon count of the fluorescence signal, which is further exacerbated by the high-speed acquisition required for capturing fast neuronal activities. The noise in voltage imaging data is often non-Gaussian and dominated by Poisson distribution, which poses a significant challenge for denoising.^7^

Deep learning-based denoising methods have shown great potential in addressing the challenges of denoising voltage imaging data. These methods have demonstrated superior performance in denoising various types of microscopy data, including fluorescence microscopy,^8^ light-sheet microscopy,^9^ and two-photon microscopy.^10^ However, in realistic denoising applications, the ground-truth high signal-to-noise ratio (SNR) measurements are often not available, which makes supervised learning-based methods less practical. In contrast, self-supervised learning-based methods have emerged as a promising alternative for denoising calcium or voltage imaging data, such as Noise2Void,^11^ Deep Interpolation,^12^ DeepVID,^1, 2^ DeepCAD (and DeepCAD-RT),^13, 14^ and SUPPORT.^15^ These self-supervised learning frameworks leverage the inherent spatial and/or temporal structure within the data, learning meaningful latent representations to perform denoising. With specifically designed tasks and loss functions, these models are adept at predicting a subset of data using the rest, bypassing the need for explicit supervision from ground truth labels. This adaptability underscores their potential for robust denoising performance in applications with limited high-SNR data availability.

In voltage imaging, achieving high spatial resolution is essential for accurately resolving fine neuronal structures, while superior temporal resolution is crucial for capturing the rapid dynamics of neuronal activities. Traditional deep learning-based methods for denoising voltage imaging data often face a significant trade-off between spatial and temporal resolution. Existing self-supervised learning frameworks typically require a large number of input frames,^12, 14^ which leads to over-smoothed temporal traces and poor temporal resolution, or they use too few frames, resulting in low spatial resolution.^1, 15^ Therefore, there is an unmet need for an advanced self-supervised denoising framework that can effectively decouple the spatial and temporal performance, and achieve superior denoising capability in resolving both fine spatial structures and rapid temporal dynamics.

In this work, we present DeepVID v2, a self-supervised denoising framework with decoupled spatiotemporal enhancement for low-photon voltage imaging. In our previous work,^1^ we introduced DeepVID, which performs frame-based voltage imaging denoising by utilizing a sequence of frames around the target frame. This leverages temporal information while ensuring reconstruction consistency. The network also integrates multiple blind pixels in the target frame to enrich learning the local spatial information. To further enhance the spatial performance, DeepVID v2 presents a novel method to preserve sharp edge information inherent in the raw data. Prior studies have demonstrated the effectiveness of utilizing edge information to improve the spatial resolution of denoised images.^16–18^ Here, by integrating an additional edge extraction branch into the DeepVID network (Fig. 1), DeepVID v2 significantly enhances the spatial resolution and integrity of the neuronal structures in the denoised images.

**Fig 1.**
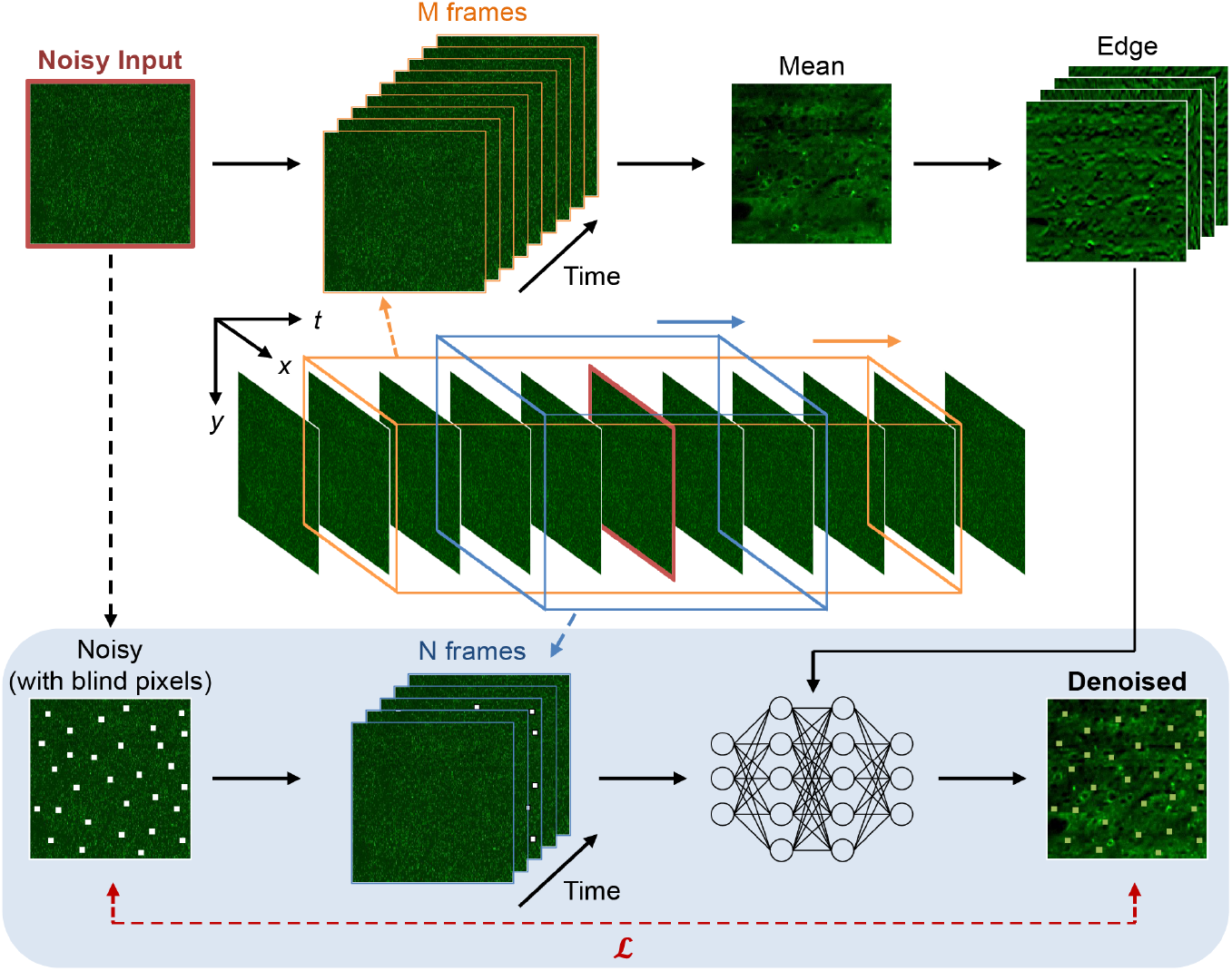
Block diagram of DeepVID v2. DeepVID v2 is composed of two main components: a main branch for denoising (bottom) and a side branch for edge extraction (top). Components adapted from our original DeepVID network are represented in the blue-shaded area.

Critically, DeepVID v2 achieves the decoupling of spatial and temporal performance by introducing two adjustable parameters: the number of input frames, *N*, and the number of frames used for edge extraction, *M*. This dual-parameter strategy enables precise fine-tuning of the denoising process, allowing for optimal resolution of both spatial structures and temporal activities, thereby overcoming the limitations observed in previous models.

We demonstrate that DeepVID v2 achieves superior spatial and temporal denoising performance under diverse imaging conditions, including various SNRs and in extreme low-photon scenarios. Our results indicate that DeepVID v2 is a promising tool for denoising *in vivo* voltage imaging data, and has the potential to facilitate the study of neuronal activities in the brain.

## 2 Methods

### 2.1 Voltage Imaging Data Collection

The data used in this study are two-photon voltage imaging image series collected from the SMURF microscope in our previous study.^1^ Spatial and temporal beam multiplexing along with a multianode photomultiplier tube (MAPMT) were used in the SMURF microscope setup. This configuration is engineered to maximize the effective repetition rate of pulsed lasers with minimal crosstalk on MAPMT, therefore enabling high-speed low-light imaging across a wide field of view (FOV). To measure the sensory-evoked neuronal responses, voltage imaging was performed at a sampling rate of 803 Hz in the primary somatosensory cortex (S1) from awake, head-fixed mice. Whisker stimulation was delivered as air puffs to the whisker pad at 10-Hz stimulus frequency in one- or five-puff trains, with 4-s intervals. The captured voltage imaging images contain 400 *×* 192 pixels in total concatenated from 8 strips (400 *×* 24 pixels per strip), with a pixel size of 1.0 *µm* along the *x*-axis and 2.1 *µm* along the *y*-axis.

### 2.2 DeepVID v2 Framework

The system diagram of DeepVID v2 is illustrated in Fig. 1. DeepVID v2 performs denoising on 3D (2D space + 1D time) image stacks on a frame-by-frame basis. The network is composed of two main components: a main branch for denoising similar to our previously developed DeepVID,^1^ and a side branch for edge extraction. DeepVID v2 utilizes both the spatial and temporal information in the raw data, as well as edge information from the side branch, to perform denoising. The neural network in the main branch is composed of four residual blocks, each containing two convolutional layers with batch normalization layers, followed by a PReLU activation layer attached after the first convolution layer. A skip connection is added between the input and output for each residual block.

Given a frame to be denoised, the side branch first takes a *M* = 2*M*_0_ + 1 image series as the input, from *M*_0_ frames before to *M*_0_ frames after the central frame, to calculate a local mean frame, resulting in an improved spatial representation than any raw single frame. A Gaussian blur filter is then applied to the mean frame to remove residual noise, followed by four Sobel filters from 0^*◦*^, 45^*◦*^, 90^*◦*^, 135^*◦*^ to extract the edge information along different directions. The outputs from the Sobel filters are then treated as four additional input channels to the main branch.

In addition to the four edge channels, the main branch takes another *N* = 2*N*_0_ + 1 image series as the input, consisting of *N*_0_ frames before and *N*_0_ frames after the central frame, as well as the degraded central frame to perform denoising. A random set of pixels are set as blind pixels in the degraded central frame with a ratio of *p*_*blind*_, whose intensities are replaced by random values sampled from the pixel intensities within the frame. These blind pixels are used to guide the network to learn the spatial and temporal information in the raw data, and to prevent the network from simply replicating the input to the output.^1^

The loss function is the mean squared error (MSE) computed between the output denoised image and the input noisy image, calculated only at the locations of the blind pixels. In this study, parameters are optimized to achieve the best spatial and temporal performance at the same time, using 7 frames as *N*, all available frames as *M*, and *p*_*blind*_ = 0.5%. The training dataset comprises 1,181 videos, with each video containing 1,000 frames captured at a rate of 803 Hz. The training utilizes the Adam optimizer with a configuration of 360 steps per epoch and a batch size of four. To avoid overfitting, the training stopped after iterating through the entire dataset three times. The initial learning rate is set to 5 *×* 10^*−*6^, then halved if the loss on the validation set plateaus over the last 288,000 samples, until it reaches the minimum learning rate of 1 *×* 10^*−*7^.

### 2.3 Spike Detection

Spike Detection is performed to infer evoked potentials from the extracted time traces. Time traces are first normalized by the mean and standard deviation of the entire time trace. For each stimulus, spikes are detected in a window from 0 s to 0.1 s after the stimulus onset, using a threshold of 3 and a minimum distance of 0.1 s between spikes. The full width at half maximum (FWHM) of the detected spikes is calculated as the time difference between the two points where the intensity reached half of the peak value. Only spikes with an FWHM falling within 3 standard deviations are retained. The number of detected spikes and the FWHM of these spikes are used to evaluate the temporal performance of the denoised videos.

### 2.4 Performance Metrics

The performance of DeepVID v2 is evaluated using a variety of metrics. Pearson Correlation Coefficient (PCC) is used to evaluate the spatial and temporal performance of DeepVID v2. The spatial PCC is determined by comparing the pixel-wise correlation between the reference frame and either the raw or DeepVID v2 denoised frame. The temporal average frame serves as the reference frame for calculating spatial PCC. Temporal PCC is computed by comparing the correlation of the reference time traces with the raw or DeepVID v2 denoised time traces. The reference time traces are obtained by employing a 7-frame moving average of the raw traces, which uses the same number of input frames as in DeepVID v2. The temporal Signal-to-Noise Ratio (SNR) is also used to evaluate the temporal performance, defined as the ratio of the mean to the standard deviation of the time trace.

### 2.5 Dataset Division based on Temporal SNR

The dataset is divided into three subsets representing low, medium, and high SNR, respectively. The averaged temporal SNR of each FOV is obtained by averaging the temporal SNR calculated from each ROI trace for all the ROIs in the FOV. The FOVs are then sorted by the averaged temporal SNR. The bottom 1/3, middle 1/3, and top 1/3 are grouped as the low, medium, and high SNR subsets, respectively.

### 2.6 Simulation of Videos in Lower-photon Regimes

In low-photon regimes, the signal is dominated by Poisson noise, in which the variance is proportional to the mean intensity. This feature is verified in our dataset by calculating the mean and variance of each single-pixel time trace (Fig. 2c). The ratio of this linear correlation, *β*, reflects the characteristic of the imaging system, and therefore should be stable in varied light conditions. The simulated video in lower-photon conditions should follow the same principle, whose variance is still proportional to the mean intensity, with the same ratio as the raw video.

**Fig 2.**
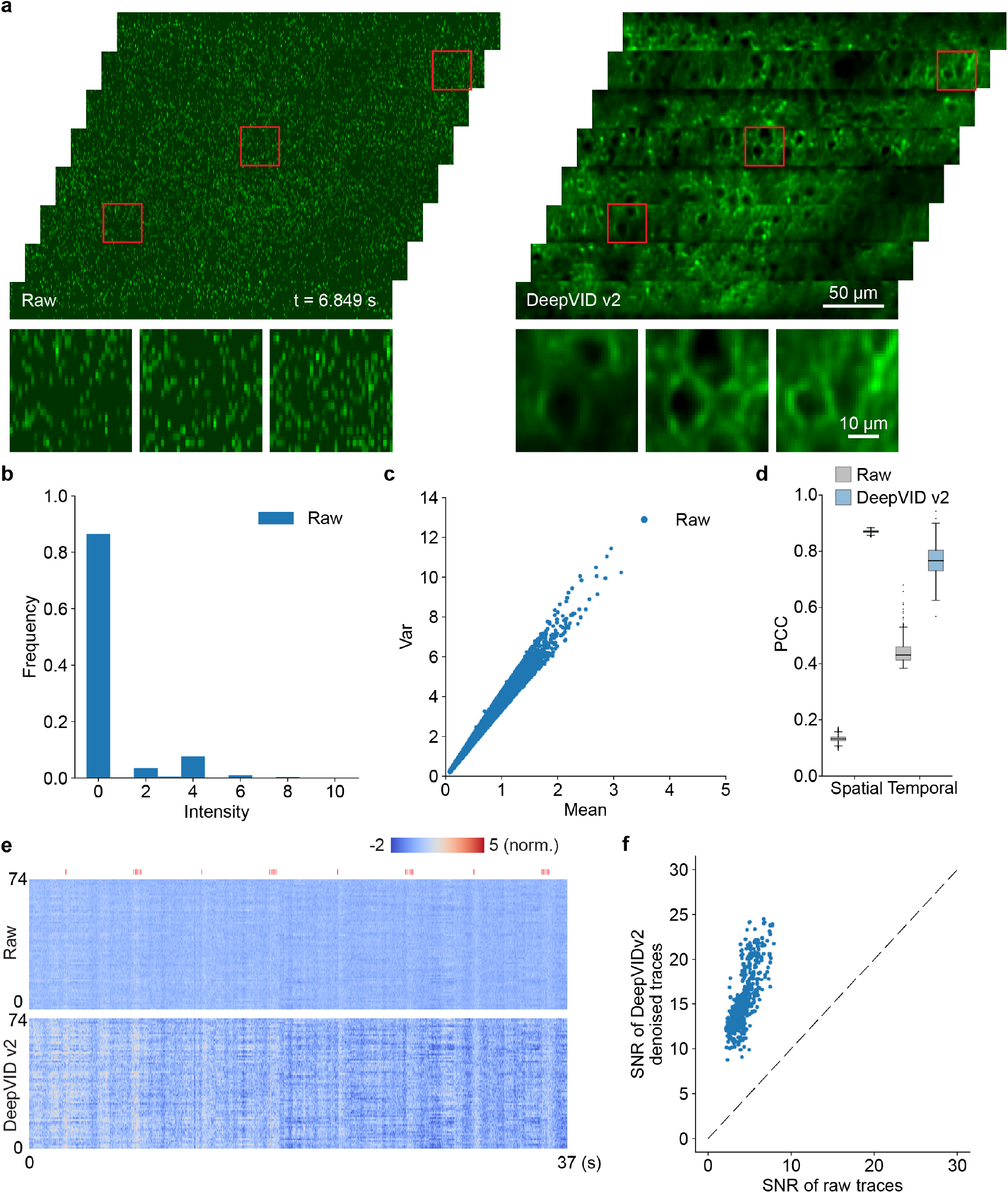
DeepVID v2 denoising enhances both the spatial and temporal quality of the voltage imaging data. (a) Single-frame images from the raw and DeepVID v2 denoised videos. (b) Histogram of the raw video. (c) Characteristics of noise in the raw video. The variance of single-pixel time traces (Y-axis) is linearly proportional to the mean of the traces (X-axis). (d) Spatial and temporal PCCs of the raw and DeepVID v2 denoised videos. (e) Heatmaps displaying time traces extracted from 74 ROIs in the raw and DeepVID v2 denoised videos. Air puff whisker stimuli are shown as red ticks on the top. (f) Temporal SNRs of the raw and DeepVID v2 denoised time traces.

To simulate voltage imaging data in lower photon regimes, we propose a two-step simulation protocol on a pixel-by-pixel basis. Before the simulation, we calculate the ratio *β*_0_ between the variance and the mean intensity for the raw video.

First, for each pixel intensity in the raw video *I*_0_, we apply Binomial degradation with a probability of *p* to obtain *I*_*b*_, and calculate the updated ratio *β*_*b*_ after applying Binomial degradation to all pixels in the video. This step reduces the intensity in the measurements, but also lowers the ratio.

Second, we multiply all pixel intensities in the simulated video *I*_*b*_ by a factor of *A* = *β*_0_*/β*_*b*_ to obtain *I*_*d*_, which increases both the intensity and the ratio by a factor of *A*. After this two-step simulation, the simulated video *I*_*d*_ has a lower intensity with a factor of *d* = *pA* compared with the raw video *I*_0_, while the ratio between the variance and the mean remains the same (Fig. S8). The proposed simulation protocol is able to simulate voltage imaging data in lower photon regimes, while maintaining the same characteristics of the imaging system.

## 3 Results

### 3.1 DeepVID v2 Improves Spatial Resolution while Preserving Temporal Dynamics

To demonstrate the denoising capability of DeepVID v2, we present single-frame full-FOV images from both the raw and DeepVID v2 denoised videos in Fig. 2a. The noisy raw video was captured in an extremely low-photon regime, with the raw pixel intensity readout lower than 10 for almost all pixels (Fig. 2b). The variance of single-pixel time traces is linear to the mean of such traces, which validates that Poisson noise dominates the raw measurements (Fig. 2c). After denoising, the membrane and other neuronal structures are clearly resolved at the single-frame level. Heatmaps displaying time traces extracted from 74 manually labeled regions of interest (ROIs) with active neurons during the 37-second measurements are depicted in Fig. 2e to highlight the improvement from the DeepVID v2 denoising. The traces from the denoised video exhibit a more pronounced contrast compared to the raw video, suggesting enhanced signals from the underlying neuronal activities after denoising.

To quantitatively assess the performance of DeepVID v2 denoising, we compute the spatial and temporal PCCs for both the raw and DeepVID v2 denoised videos, illustrated in Fig. 2d. Both the spatial and temporal PCC values of the denoised video significantly surpass those of the raw video (spatial PCC: 0.90 *±* 0.01 for DeepVID v2, 0.15 *±* 0.01 for raw, *n* = 38972; temporal PCC: 0.77*±* 0.06 for DeepVID v2, 0.44*±* 0.04 for raw, *n* = 516), indicating that DeepVID v2 effectively denoises the raw voltage imaging video in both the spatial and temporal domains.

Furthermore, we calculate the temporal SNRs for the raw and DeepVID v2 denoised time traces extracted from all ROIs in the FOV, as presented in Fig. 2f. The temporal SNRs of denoised time traces are consistently higher than that of the raw traces for all ROIs (DeepVID v2, 15.57 *±* 3.19; raw, 4.25 *±* 1.17; *n* = 516), further underscoring the effective temporal denoising capability of DeepVID v2.

Next, we investigate the performance of DeepVID v2 with a focus on single-neuron activities. From another time-series measurement with a single-frame full-FOV denoised image shown in Fig. 3a, we extract a few key frames from an ROI in Fig. 3d, along with the corresponding raw frames in Fig. 3b. The time traces from an active neuron (circled in red in Fig. 3d) are extracted from both the raw and denoised videos, as shown in Fig. 3c and Fig. 3e, respectively. The activation on the neuronal membranes is consistently resolved in the zoom-in images at the timestamps marked on the time traces.

**Fig 3.**
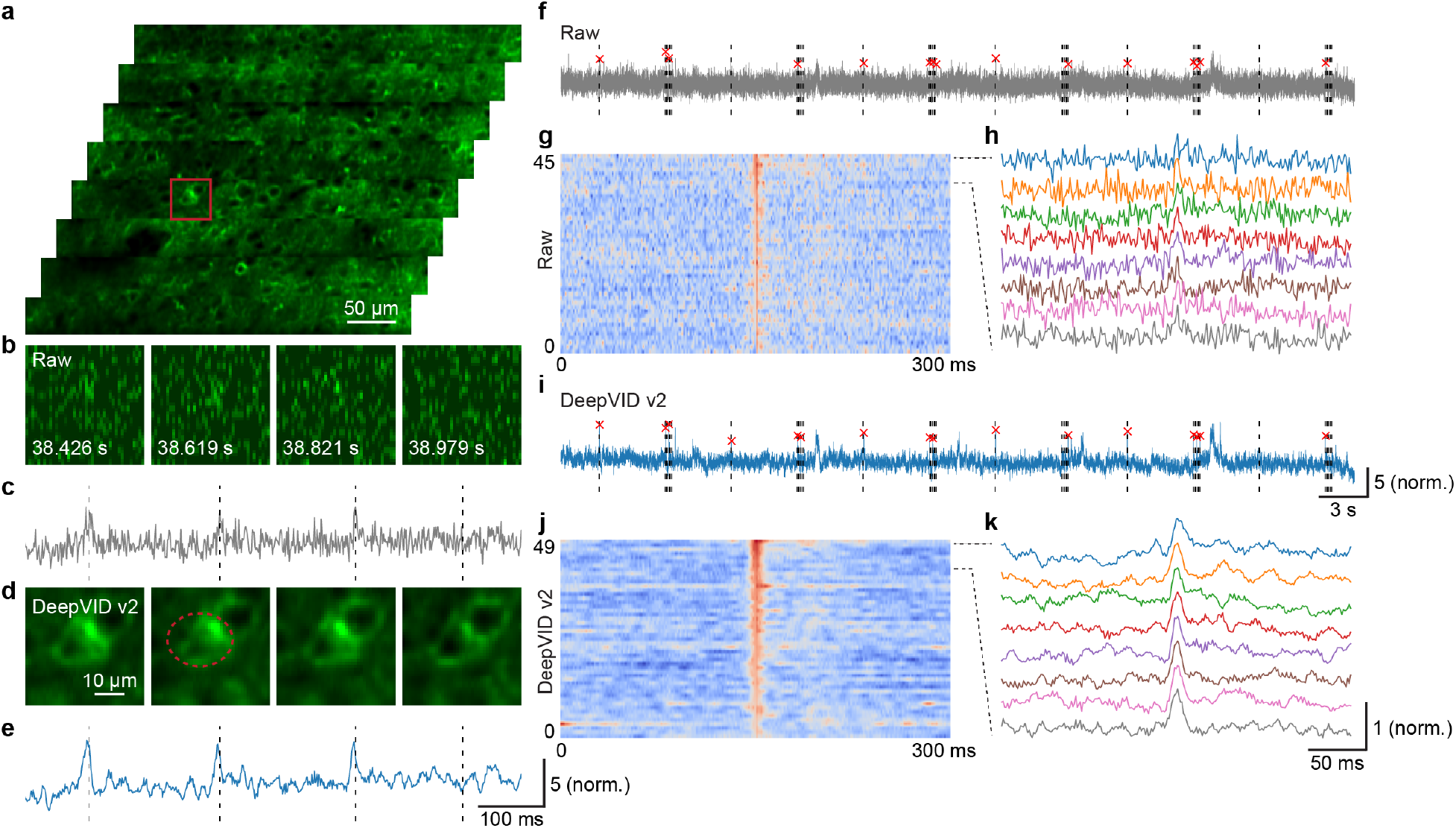
Denoising performance on single-neuron activities. (a) A single-frame full-FOV denoised image. (b) Zoom-in view and (c) time trace of the ROI from the raw video. (d) Zoom-in view and (e) time trace of the ROI from the DeepVID v2 denoised video. (f) Detected evoked potentials, (g) heatmap of the detected evoked potentials, and (h) time traces of the detected evoked potentials from the raw video. (i) Detected evoked potentials, (j) heatmap of the detected evoked potentials, and (k) time traces of the detected evoked potentials from the DeepVID v2 denoised video. Air puff whisker stimuli are shown as dotted lines in (f) and (i).

We further apply spike detection on the time traces to extract evoked potentials (Fig. S1). The evoked potentials extracted are marked in red, while the stimuli are shown as dotted lines in Fig. 3f and Fig. 3i. All 45 detected evoked potentials are aligned at the peak and presented as heatmaps in Fig. 3g and Fig. 3j. The time traces of the evoked potential are displayed in Fig. 3h and Fig. 3k. The evoked potentials extracted from the denoised video exhibit less noisy traces compared to the raw video, which indicates the improved capability of DeepVID v2 in resolving neuronal activities.

### 3.2 DeepVID v2 Overcomes Tradeoff Between Spatial and Temporal Performance

The performance of previous self-supervised denoising algorithms has often been influenced by the tradeoff between spatial and temporal resolution, which is controlled by the number of input frames *N* to the network (Fig. S5 and Fig. S6). Unlike previous methods, DeepVID v2 is designed to decouple spatial and temporal performance by incorporating two key parameters: the number of input frames, *N*, and the number of frames used for edge extraction, *M*. To investigate the effect of these parameters on the performance of DeepVID v2, we vary *N* and *M* and train a neural network model for each combination.

First, we fix *M* as the maximum available frames and vary *N* from 3 to 127. As *N* increases, time traces become over-smoothed and spikes become harder to recognize, as shown in Fig. 4a and Fig. S2. To evaluate the temporal performance, spike detection is performed on the time traces, and the temporal metrics including the number of detected spikes and the FWHM of the detected spikes are calculated. The FWHM of the detected spikes increases with increasing *N*, while the number of detected spikes initially increased due to improved temporal SNR but later decreased as the traces become over-smoothed, as shown in Fig. 4b. The spatial PCCs of the denoised videos are significantly higher than that of the raw video but remain comparable across different *N*, indicating that the spatial performance is not significantly affected by *N*, as shown in Fig. 4c and Fig. S3. From this analysis, we conclude that the optimal value for *N* is 7 for our experimental conditions, which provides the best combination of narrow FWHMs and a large number of reliably detected spikes.

**Fig 4.**
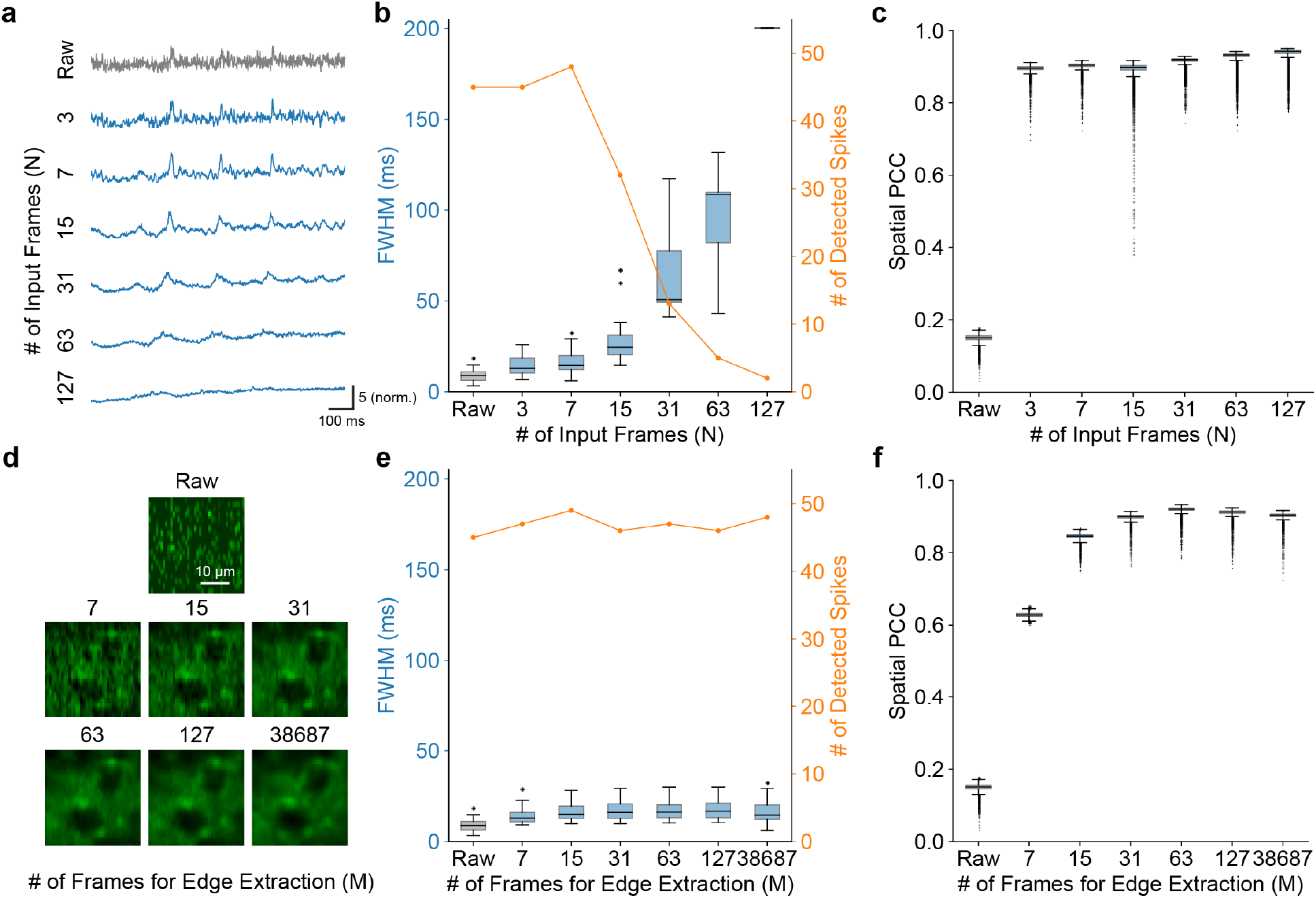
Parameter analysis. (a) Time traces extracted from the same ROI from the DeepVID v2 denoised videos with different *N*. (b) Temporal metrics and (c) spatial PCC of the DeepVID v2 denoised videos with different *N*. (d) Zoom-in view of a ROI from a single-frame image in the DeepVID v2 denoised videos with different *M*. (e) Temporal metrics and (f) spatial PCC of the DeepVID v2 denoised videos with different *M*.

Next, we fix *N* at 7 frames for the optimal temporal performance and vary *M* from 7 to the maximum available frames. Fig. 4d shows the zoomed-in views of an ROI from a single-frame image in the raw and denoised videos using DeepVID v2 with different *M* indicated above each image. The number of detected spikes and the FWHM of the detected spikes are comparable across different *M*, indicating that the temporal performance is not significantly affected by *M*, as shown in Fig. 4e and Fig. S4. The spatial PCCs of the denoised videos increase with *M*, suggesting better spatial performance with an increased *M*, as shown in Fig. 4f.

Our parameter analysis reveals that the new framework of DeepVID v2 is able to decouple the spatial and temporal performance by independently adjusting *M* and *N*. This decoupling enables DeepVID v2 to achieve superior denoising capability in resolving both spatial neuronal structures and temporal neuronal activities with high SNR.

We further conduct a comparative analysis of DeepVID v2 against other recently developed self-supervised denoising methods, including Deep Interpolation,^12^ SUPPORT,^15^ DeepCAD-RT,^14^ and our previously developed DeepVID^1, 2^. All benchmarks, except DeepCAD-RT, utilize *N* = 7 frames as input to the network, whereas DeepCAD-RT adheres to the default optimal setting of *N* = 127 after splitting odd and even stacks, based on our own extensive parameter search. DeepVID v2 uses all available frames (*M*) for edge extraction. Our evaluation focuses on spatial performance using single-frame images and temporal performance using time traces extracted from ROIs (Fig. 5a). Regarding the temporal performance, we conduct spike detection for each benchmark. DeepVID v2 demonstrates strong temporal performance in terms of the number of detected spikes and FWHM of the detected spikes (Fig. 5b), in contrast to the over-smoothed time traces observed in DeepCAD-RT with *N* = 127. DeepVID v2 also exhibits superior spatial performance in terms of spatial PCC (Fig. 5c), matching the performance of DeepCAD-RT using many more input frames, and surpassing other benchmarks.

**Fig 5.**
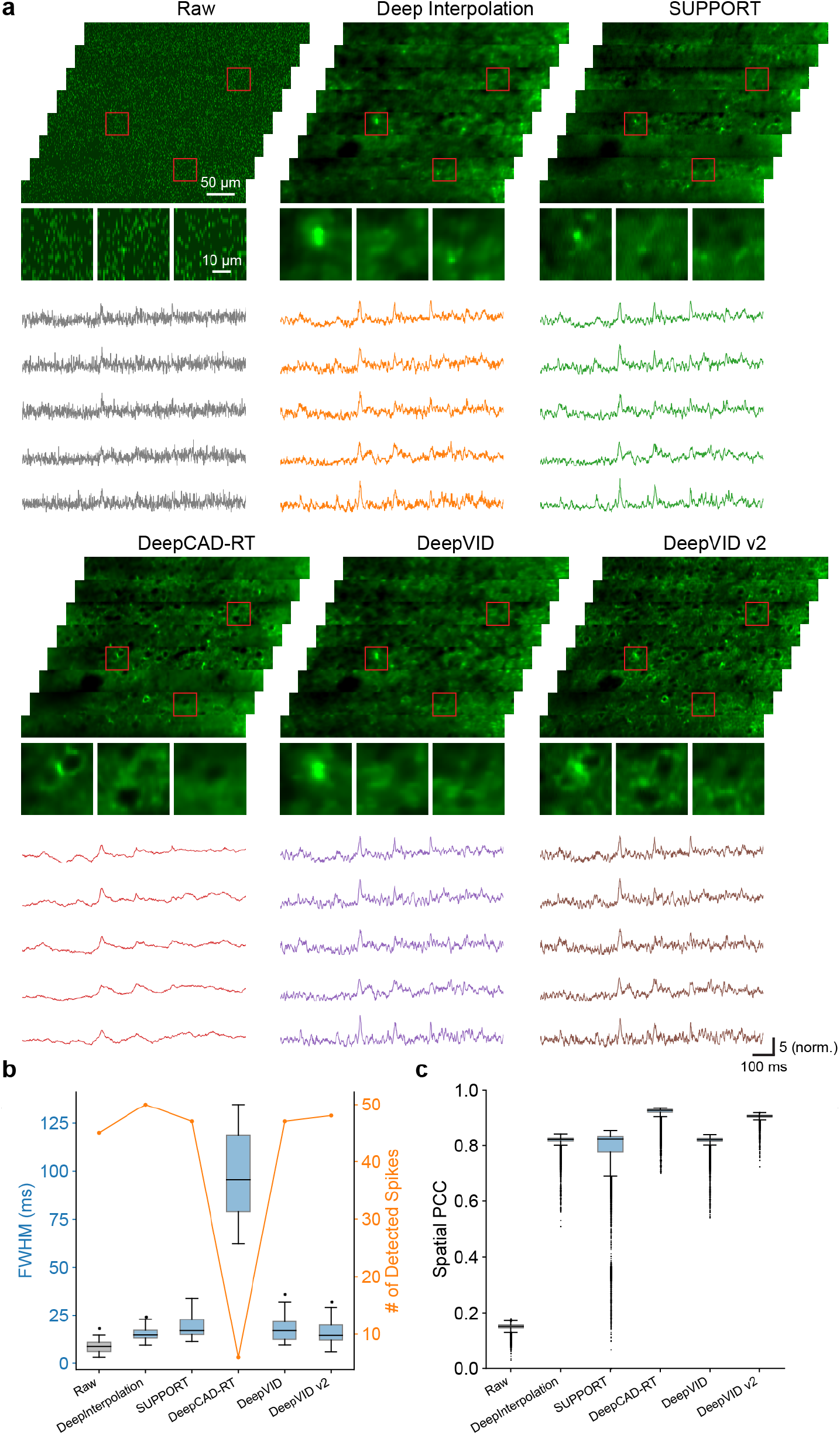
Benchmark comparison. (a) Single-frame images and ROI time traces from the raw and denoised videos. (b) The number of detected spikes and FWHM of the detected spikes from the raw and denoised ROI time traces. (c) Spatial PCC of the raw and denoised videos.

To characterize the benchmark performance given similar inputs, we also compare the performance of all benchmark networks in two other conditions, including one with a small *N* at 7 frames (Fig. S5), and another with a large *N* at 127 frames (Fig. S6). DeepVID v2 maintains the same optimal parameter settings, with *N* = 7 and *M* using all available frames. When *N* is small at 7 frames, all benchmarks show similar temporal performance in terms of small FWHM of detected spikes, while jagged edges are observed only in DeepCAD-RT, as limited frames are available for 3-D convolution in the time axis. SUPPORT shows good but unstable spatial performance, with clear structures in some high spatial SNR strips, but with a strong blur in other lower spatial SNR strips. When *N* is large at 127 frames, all benchmarks show similar spatial performance with high spatial PCC, but only DeepVID v2 maintains the optimal temporal performance with small FWHM of detected spikes.

Except for DeepVID v2, all other benchmarks encounter a trade-off between spatial and temporal performance: achieving good spatial but poor temporal performance with small *N* (Fig. S5), and vice versa with large *N* (Fig. S6). By decoupling spatial and temporal performance into two parameters, DeepVID v2 successfully overcomes this trade-off, thereby achieving superior performance in both spatial and temporal metrics simultaneously.

### 3.3 DeepVID v2 Generalizes to Different Imaging Conditions

To evaluate the generalization capability of DeepVID v2 under different imaging conditions, we test apply it to voltage imaging data with various SNRs. To perform this study, our experimental dataset is divided into three subsets, labeled as low, medium, and high SNR groups. The data division is based on the averaged temporal SNR calculated on all the manually labeled ROIs for each FOV in our dataset, as detailed in Sec. 2.5. A separate DeepVID v2 model is trained for each subset. The temporal SNR of the time trace for each ROI after denoising using each model is calculated and presented in Fig. 6a. The ROIs are sorted by the temporal SNR of the raw time traces, which spans a wide range from 1.14 to 10.17, as shown in the black curve. The temporal SNR after denoising by all three DeepVID v2 models consistently improves by over 3.2 fold for all ROIs (low SNR model, 3.20 *±* 0.86; medium SNR model, 3.26 *±* 0.87; high SNR model, 3.28 *±* 0.90; *n* = 2717), regardless of the SNR of the raw video. The spatial (Fig. 6b) and temporal PCC (Fig. 6c) of the denoised videos are significantly higher than that of the raw video, and importantly remain consistent across all models trained with different SNRs (see Fig. S7 for single-frame images and time traces), indicating that DeepVID v2 is able to generalize to data with various SNRs.

**Fig 6.**
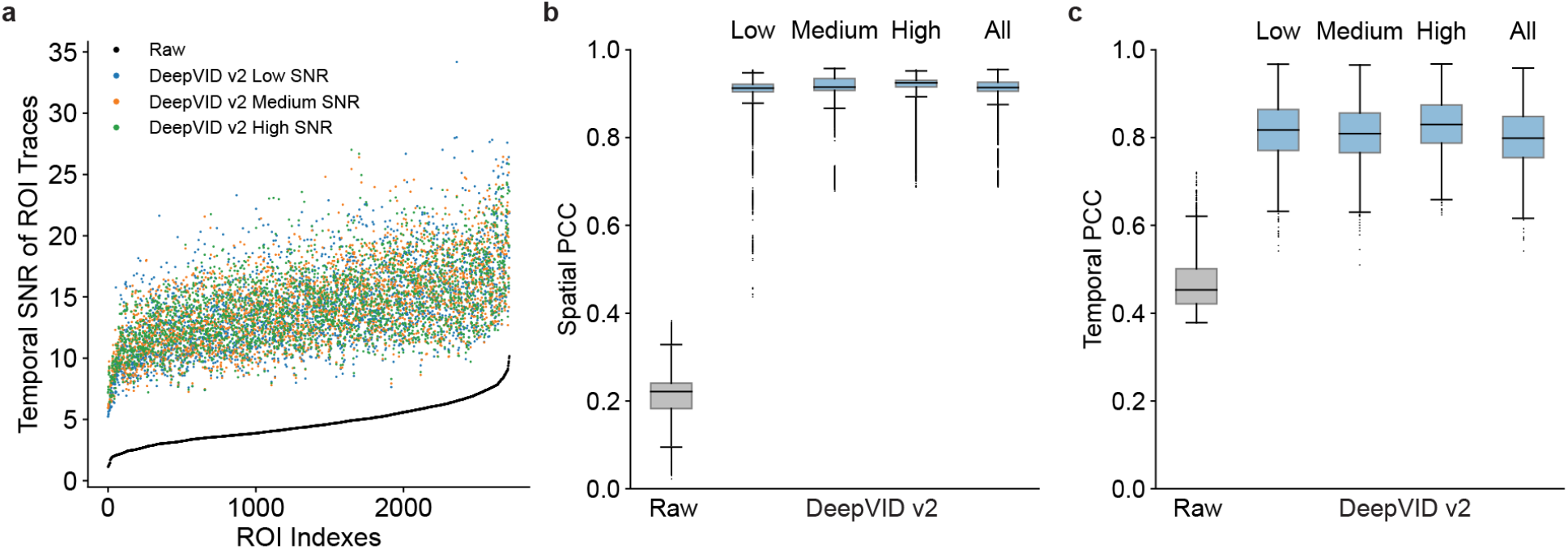
Generalization over measurements with various SNRs. (a) Temporal SNR of time traces from the raw measurements and DeepVID v2 trained using data with different SNRs. (b) Spatial PCC and (c) temporal PCC of the raw and DeepVID v2 denoised videos.

We further evaluate DeepVID v2 in extreme low-photon regimes. To perform this study, we simulate voltage imaging data at various photon levels from 10% to 100% with respect to the original measurement (see details in Sec. 2.6 and Fig. S8), and test the denoising performance of DeepVID v2. For each photon level, we train a separate DeepVID v2 model, which is possible as DeepVID v2 is a self-supervised method that does not need external ground-truth data for training. The zoom-in image of an ROI and the time traces extracted from the simulated noisier measurements and the denoised videos are presented in Fig. 7a and Fig. 7b, respectively. DeepVID v2 is able to reliably perform denoising on data with photon levels as low as 30% of the original measurement. Both the temporal PCC (Fig. 7c) and spatial PCC (Fig. 7d) show dramatic improvements after denoising for photon levels as low as 30% of the original. The results underscore DeepVID v2’s ability to perform voltage imaging denoising in extreme low-photon conditions, which is critical for further pushing the limits in voltage imaging *in vivo*.

**Fig 7.**
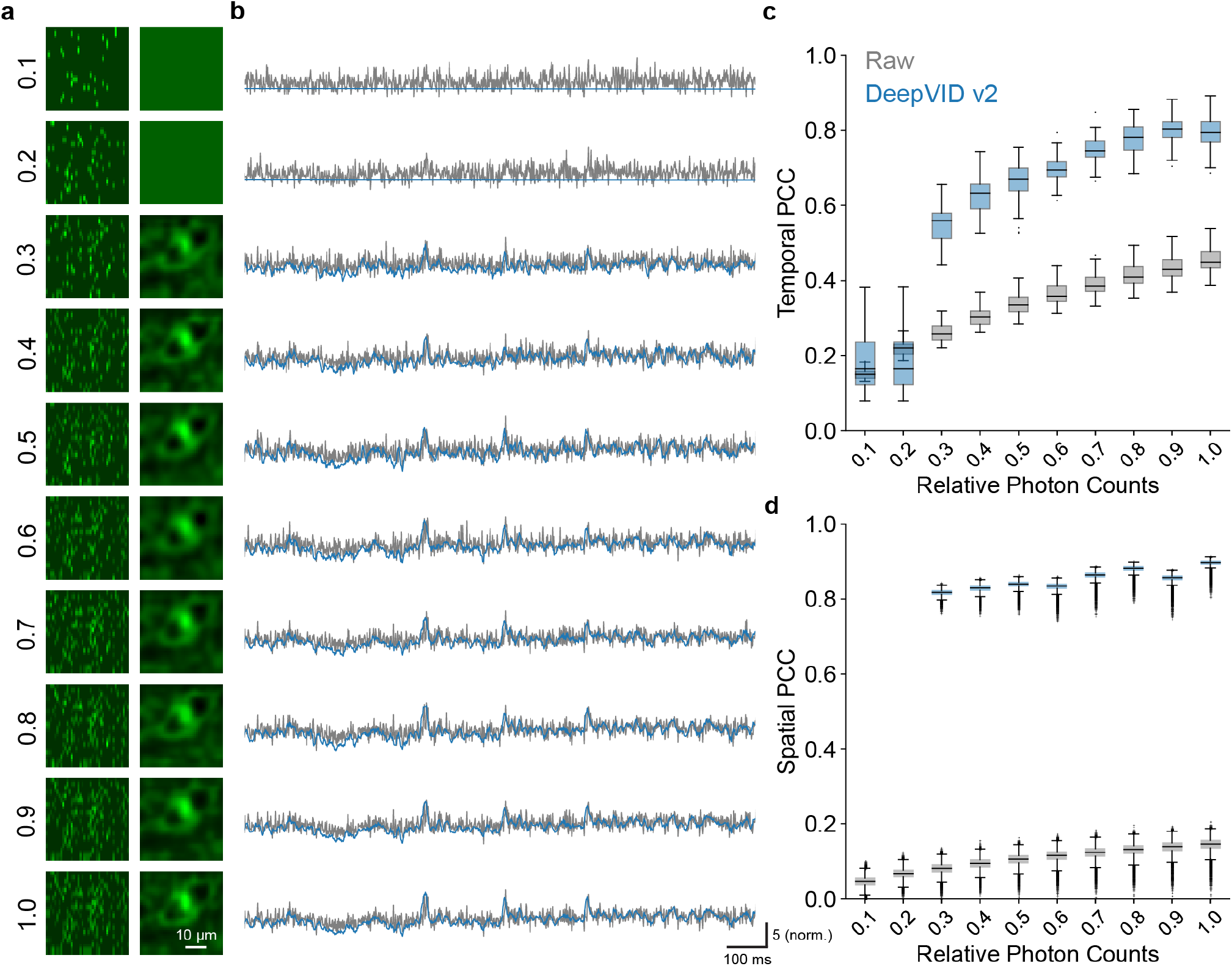
Simulation of DeepVID v2 denoising capability in extreme low-photon regimes. (a) Zoom-in image of an ROI and (b) the time traces from the simulated noisy and denoised videos. (c) Temporal PCC of the simulated noisy and denoised time traces. (d) Spatial PCC of the simulated noisy and denoised videos.

## 4 Discussion

We introduced DeepVID v2, a self-supervised denoising framework with decoupled spatiotemporal enhancement capability for low-photon voltage imaging. By integrating an additional edge extraction branch into the DeepVID architecture^1^ with two adjustable parameters, DeepVID v2 effectively addresses the inherent tradeoff between spatial and temporal performance, resulting in enhanced denoising capabilities for resolving both spatial neuronal structures and temporal dynamics. Additionally, our experiments demonstrated the robustness of DeepVID v2 across diverse imaging conditions, including videos with varying SNR and measurements simulated under extreme low-photon regimes. These results highlight DeepVID v2’s potential as a valuable tool for denoising voltage imaging data, offering promising prospects for advancing the study of neuronal activities within the brain.

As a limitation of this work, the performance of DeepVID v2 may be influenced by the relationship between imaging speed and the object’s motion. The extraction of edge information in DeepVID v2 is based on the assumption that the local mean frame is clean without motion artifacts, which may not hold true when the moving speed of the object is much faster than the imaging speed. In such scenarios, the framework used in SUPPORT would be an alternative solution as it demonstrated its denoising capability on moving *C.elegans* when the object’s locomotion was faster than the imaging speed. In this case, spatial information from neighboring pixels in the central frame contributes more to the denoising process in SUPPORT, rather than temporal information from adjacent frames.^15^ However, the spatial performance of the SUPPORT denoised result may be sensitive to the spatial quality of the raw measurements, as observed in the varying spatial performance for different strips in our benchmark comparison (Fig. 5). It remains a challenge to achieve robust denoising performance in high-speed low-photon large-FOV imaging scenarios where object movement surpasses the imaging speed, indicating the need for further methodological developments to address this issue effectively.

### Disclosures

The authors declare no conflicts of interest.

### Code, Data, and Materials Availability

Code for PyTorch implementation of DeepVID v2 is available at the GitHub repository: https://github.com/bu-cisl/DeepVIDv2. Data used in this study are obtained from a previous publication.^1^

## Acknowledgments

This work was supported by the National Institutes of Health (NIH) under grant U01NS128665.

## Notes

https://github.com/bu-cisl/DeepVIDv2

### Competing Interest Statement

The authors have declared no competing interest.

